# Chemogenetic Breakdown of the Dentate Gate Causes Seizures and Spatial Memory Deficits

**DOI:** 10.1101/2024.11.12.623184

**Authors:** Christopher D. Adam, Emily D. Schellinger, Alicia White, Srdjan M. Joksimovic, Hajime Takano, Douglas A. Coulter

## Abstract

The dentate gyrus has often been posited to act as a gate that dampens highly active afferent input into the hippocampus. Effective gating is thought to prevent seizure initiation and propagation in the hippocampus and support learning and memory processes. Pathological changes to DG circuitry that occur in temporal lobe epilepsy (TLE) can increase DG excitability and impair its gating ability which can contribute to seizures and cognitive deficits. There is evidence that TLE pathologies and seizures may independently contribute to learning and memory deficits in TLE through distinct mechanisms. These two factors are difficult to untangle since TLE pathologies can drive seizures, and seizures can worsen TLE pathologies. Here we assessed whether chemogenetically increasing dentate granule cell (DGC) excitability was enough to break down the dentate gate in the absence of TLE pathologies. We found that increasing excitability specifically in DGCs caused seizures in non-epileptic mice. Importantly, due to the modulatory nature of DREADD effects, seizures were driven by intrinsic circuit activity rather than direct activation of DGCs. These seizures resulted in a spatial memory deficit when induced after training in the spatial object recognition task and showed stereotypical patterns of activity in miniscope calcium recordings. Our results provide direct support for the dentate gate hypothesis since seizures could be induced in non-epileptic animals by artificially degrading the dentate gate with chemogenetics in the absence of epilepsy pathologies.

## INTRODUCTION

Epilepsy affects approximately 1% of the population worldwide making it one of the most common neurological disorders in the world [1]. Temporal lobe epilepsy (TLE) is the most prevalent clinical variant of epilepsy in adults and is associated with structural and functional changes to neuronal circuits in the temporal lobe. In addition to the spontaneous recurring seizures that define all epilepsies, TLE patients commonly present with cognitive comorbidities including memory impairment, anxiety, and depression, and they report these cognitive comorbidities, and the stigmas associated with them, to be severely debilitating and significantly decrease their quality of life [2–8].

The hippocampal dentate gyrus (DG) is implicated in TLE where it has been posited to act as a ‘gate’, preventing excessive activity in the hippocampus [9, 10]. This gating function is important for cognitive processes but also prevents pathological discharges in the hippocampus that can be epileptogenic. Dentate gating is supported by the low excitability and sparse firing of dentate granule cells (DGCs), the primary output cells of the DG, and is driven by DGC intrinsic properties and DG circuit properties. The especially hyperpolarized resting membrane potential and low input resistance of DGCs results in a relatively high threshold for activation [11]. Additionally, DGCs receive strong feedforward and feedback inhibition from numerous interneuron populations including dendritic inhibition in the molecular layer and strong shunting inhibition in the granule cell layer [12, 13]. DGCs also outnumber their EC inputs by a factor of about 5:1 [14], and this large expansion ratio allows promiscuously firing EC inputs to be orthogonalized across the large DGC network implicating the DG in pattern separation computations. The summed result of these intrinsic and circuit properties is a dentate gate that supports learning and memory processes and helps to prevent seizure initiation and propagation throughout the hippocampus.

In TLE, pathological changes in DG circuitry cause DGCs to become hyperexcitable which contributes to the breakdown of the dentate gate [9, 10]. This process is thought to make the hippocampus susceptible to seizures and directly disrupt learning and memory functions supported by the DG. Pathological changes include a loss of DG interneurons [15] and mossy cells [16–26], changes in DGC GABA receptor expression [27, 28], mossy fiber sprouting [29], and increased neurogenesis and ectopic expression of DGCs in the hilus [30]. All of these pathologies sum to massively increase circuit excitability in the DG through both intrinsic and synaptic mechanisms [31–34]. This degrades its gating ability resulting in an increased susceptibility to seizures and disrupted cognition.

An important study found that decreasing DG excitability in epileptic mice was able to rescue spatial memory deficits, while increasing DG excitability in controls mimicked behavioral deficits seen in TLE mice [35]. This supports the idea that DG hyperexcitability, driven by TLE pathologies, can affect learning and memory. Decreasing DG excitability had no effect on seizures in this study, and mice were monitored during and proximal to behavior to ensure seizures did not occur around behavioral testing. This is important because seizures can exacerbate cognitive deficits through retrograde amnesia and confusion during postictal depression. Disentangling how TLE pathologies and seizures independently affect learning and memory is difficult since seizures can cause circuit pathologies, and pathologies can lead to seizures. Studies specifically investigating how seizures affect cognition found that optogenetically shortening seizure durations in epileptic mice can improve spatial memory [36] and inducing seizures in non-epileptic mice with PTZ can disrupt spatial memory [37]. While these studies support a specific role of seizures in cognitive deficits, the role of the DG in this process is unclear. One study found that optogenetically activating DGCs can cause seizures in non-epileptic mice [38]; however, behavioral effects of this stimulation have not been investigated. Additionally, optogenetic stimulation results in direct and artificial activation of DGCs. In this study we assessed whether intrinsic circuit activity would drive seizures when DGCs were made to by hyperexcitable. We found that chemogenetically increasing DGC excitability caused seizures in non-epileptic mice. Additionally, mice showed spatial memory deficits when seizures were induced just after learning. These results provide direct evidance for the dentate gate theory and demonstrate that TLE-relevant seizures can disrupt learning and memory processes in the absence of TLE pathologies.

## RESULTS

### Increasing excitability specifically in DGCs causes secondary generalized seizures in non-epileptic mice

The dentate gyrus (DG) has been hypothesized to act as a gate to dampen highly active afferent input into the hippocampus, and breakdown of this gate is thought to contribute to seizure initiation and propagation in temporal lobe epilepsy (TLE) [9, 10, 39]. A previous study showed that optogenetically driving dentate granule cells (DGCs) to fire caused seizures in non-epileptic mice [38]; however, this artificial stimulation paradigm did not assess whether intrinsic inputs could drive seizures when the dentate gating function was compromised. To assess this possibility, we examined whether DREADD-mediated excitation of DGCs would break down the dentate gate and cause seizures in non-epileptic mice. This strategy allowed us to increase excitability specifically in DGCs and investigate whether intrinsic circuit activity would drive seizures when DGCs were made to be hyperexcitable. This was achieved by injecting a cre-specific virus containing the excitatory DREADD hM3Dq into the dentate gyrus of Rbp4-cre mice. These mice express cre recombinase in layer V cortical neurons and DGCs, but targeted injections resulted in specific expression of hM3Dq in DGCs (**Figure 1A**) allowing for specific modulation of DGC excitability. Following viral injections, mice were implanted with a twisted wire electrode in the hippocampus and 2 cortical screws (**Figure 1A**), then underwent continuous video-EEG monitoring to look for seizures. After a baseline recording period of 1 week, mice were given a saline injection followed by 4 different doses of CNO (0.1, 0.3, 1, and 3 mg/kg). CNO injections were given in a random order with a minimum washout of 48 hours between the 0.1, 0.3, and 1 mg/kg doses and a minimum washout of 72 hours after the 3 mg/kg CNO dose. EEG traces were manually reviewed *post hoc* for secondary generalized seizures that occurred in both the hippocampus and cortex simultaneously.

**Figure 1:**
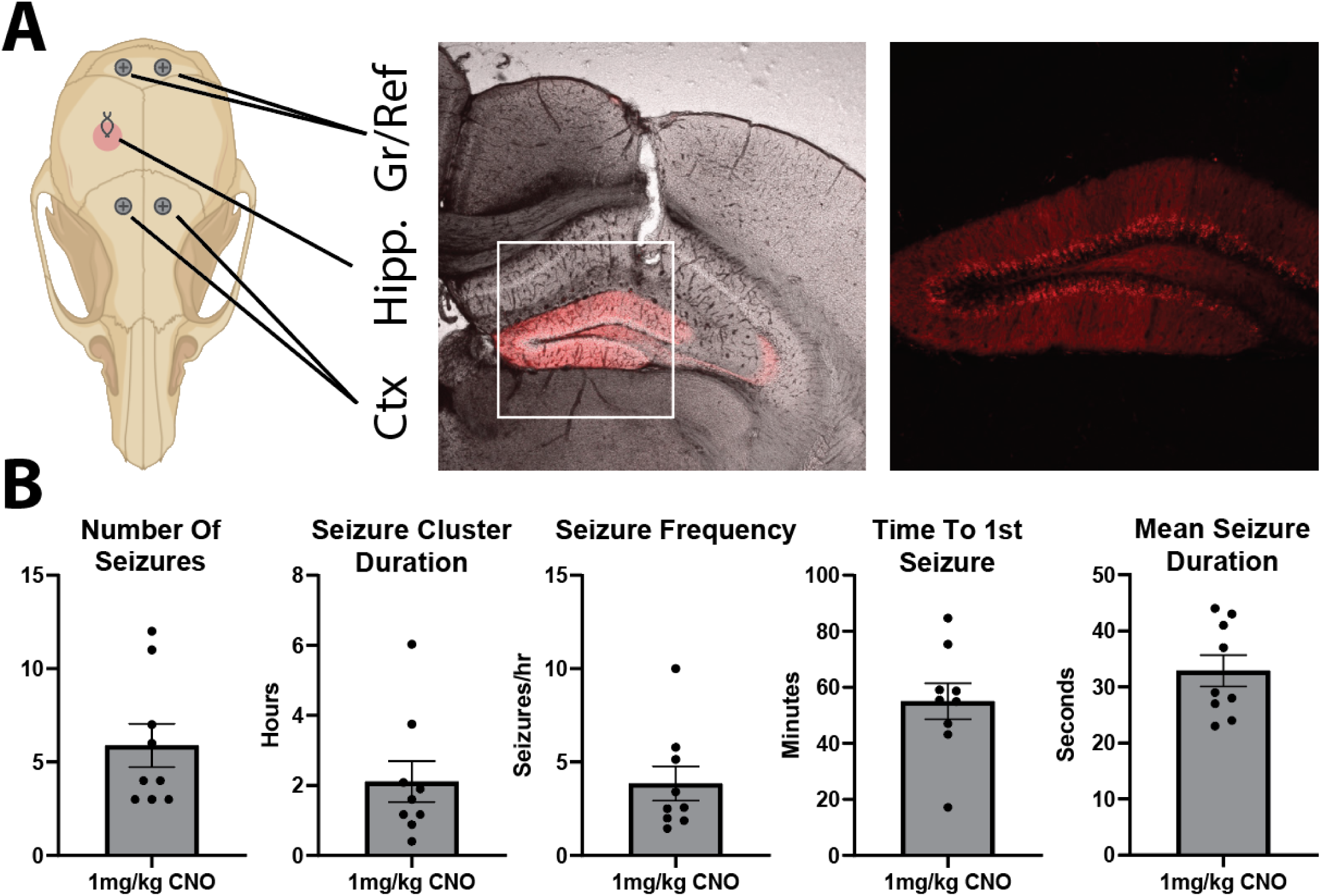
Increasing excitability specifically in DGCs reliably induces seizures. **A:** Schematic of the EEG recording electrodes and representative confocal images showing the hippocampal electrode track and hM3Dq-mCherry expression. Zoomed in fluorescence only image demonstrates that hM3Dq expression is specific to DGCs. **B:** Summary figures of seizure outcome measures following 1 mg/kg CNO injection (n = 9 mice; number of seizures: 5.89 ± 1.16; seizure cluster duration: 2.11 ± 0.58 hrs; number of seizures per hour: 3.86 ± 0.91; time to first seizure: 55.07 ± 6.41 mins; average seizure duration: 32.89 ± 2.79 seconds; all mean ± SEM).

No secondary generalized seizures occurred in mice that received 0.1 or 0.3 mg/kg CNO. However, these mice occasionally showed small hippocampal seizures as well as interictal spikes within the hippocampus or across the hippocampus and cortex. At 1 mg/kg CNO, 90% (9/10) of mice had secondary generalized seizures. These seizures occurred in a cluster of 3-12 seizures (5.89 ± 3.48; mean ± StDev) over a period of 0.40-6.03 hours (2.11 ± 1.75; mean ± StDev) resulting in an average of 3.86 ± 2.74 seizures an hour (mean ± StDev; **Figure 1B**). The average time to the first seizure in the cluster was 55.07 ± 19.23 minutes (mean ± StDev), and the average duration of each seizure in the cluster was 32.89 ± 8.36 sec (mean ± StDev; **Figure 1B**).

**Figure 2** shows a representative secondary generalized seizure. Seizures were sometimes visible on the hippocampal electrodes just before the cortical electrodes, while other times, seizures appeared on both simultaneously. Some animals also had hippocampal seizures that did not secondarily generalize. These events were especially apparent in the one animal that did not have any secondary generalized seizures. Mice often had interictal spikes either preceding or following secondary generalized seizures, and these spikes were sometimes specific to the hippocampus but frequently occurred in both the hippocampus and cortex simultaneously. When interictal spikes were visualized in both the hippocampus and cortex, spikes occurred on both sides of the cortex simultaneously.

**Figure 2:**
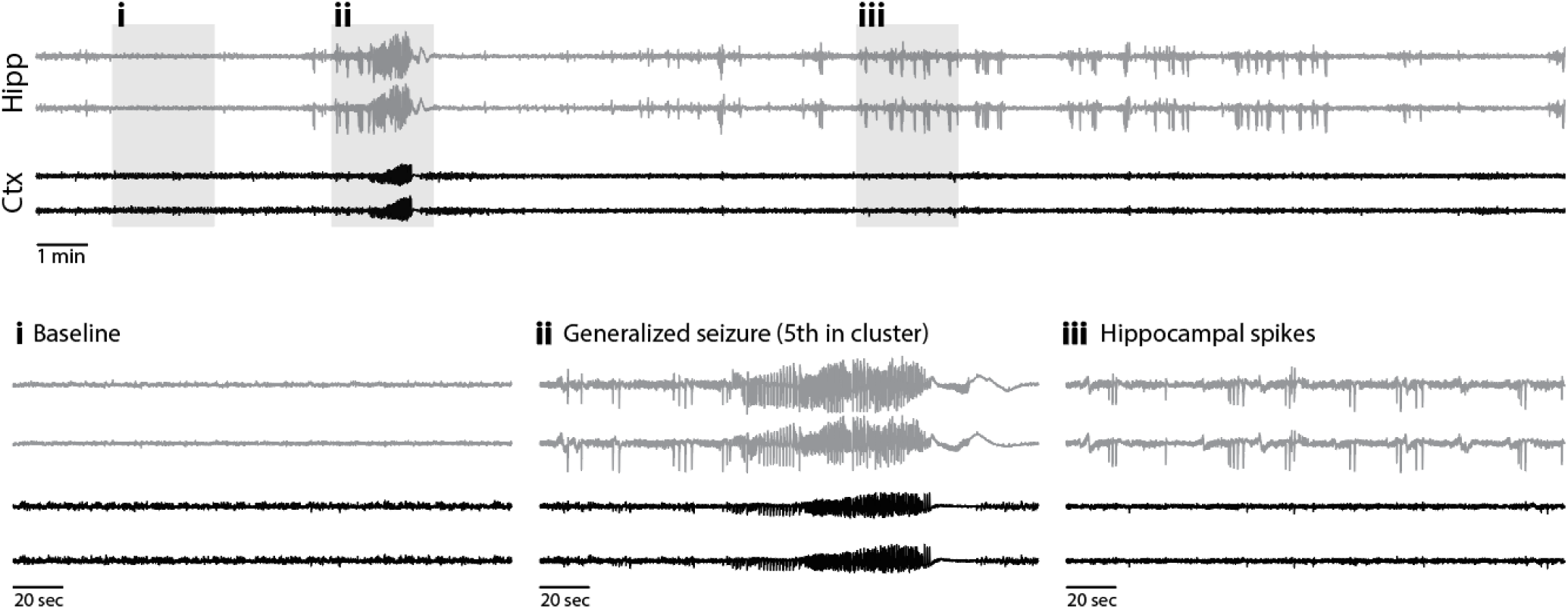
Representative seizure traces. **Top:** 30 min trace from a mouse 108 mins after 3 mg/kg CNO injection. **Bottom:** 2-minute segments (gray boxes) expanded below show: (i) baseline activity, (ii) secondary generalized seizure, & (iii) hippocampal spikes.

At 3 mg/kg CNO, all 10 mice had secondary generalized seizures, but individual animal responses became more heterogeneous. Secondary generalized seizures were more difficult to quantify due to long periods of interictal spikes and high frequency synchronization across the hippocampus and cortex that could last tens of minutes to hours. These periods often had a slow ramp up and/or down which made it difficult to define their starting and stopping point in a consistent manner across animals. In addition, some animals had long periods of hippocampal seizures in the absence of cortical seizures or in the presence of cortical spikes. These events, which could last tens of minutes, often preceded secondary generalized seizures. Some animals also had long periods of postictal depression during which hippocampal and cortical electrodes were often coherent or had coherent spiking events. Overall, the 3 mg/kg CNO dose led to more heterogenous results that made comparisons across animals and doses difficult; however, the seizure burden at this dose was higher than at 1 mg/kg.

An additional 3 animals received viral injections and underwent EEG monitoring but were excluded from the above seizure analyses. Two mice had no expression of DREADDs anywhere in the brain and were likely rbp4-cre negative mice that were incorrectly genotyped. Both of these mice did not exhibit seizures at 3 mg/kg CNO. Additionally, 1 of the 2 mice was tested at 10 mg/kg CNO and did not exhibit seizures. This, paired with the lack of generalized seizures in all mice at baseline or following the saline injection, suggests that seizures were in fact elicited by specific CNO-mediated hM3Dq activation and not from non-specific effects of CNO administration or from any potential constitutive activity of hM3Dq (known to occur in G-protein coupled receptors). The third mouse that was excluded only had unilateral expression of hM3Dq on the side of the hippocampal electrode. This mouse did not have any secondary generalized seizures at 1 mg/kg CNO; however, small hippocampal events and interictal spikes were observed. Interestingly, this mouse did have secondary generalized seizures when given 3 mg/kg CNO.

The CNO dose dependence of seizure severity suggests that dentate gate breakdown is graded with small increases in DGC excitability resulting in focal hippocampal seizures and larger increases in DGC excitability resulting in secondary generalized seizures. Importantly, secondary generalized seizures were reliably induced at a low to moderate dose of CNO (1 mg/kg) and were relatively homogenous in their presentation especially when compared to seizures induced by 3 mg/kg CNO administration.

### Induced seizures impair spatial memory

DG hyperexcitability has been demonstrated in TLE and is thought to contribute to learning and memory deficits independent of seizures [31–35, 40]. However, the effect of DG-driven seizures on learning and memory in the absence of TLE pathologies is unknown especially when testing occurs outside of the postictal depression period. To assess this, we investigated whether DREADD-induced seizures could impair spatial memory in non-epileptic mice. In order to verify seizure activity and investigate seizure effects on coding, we performed calcium imaging from freely behaving mice using miniscopes. This was done through viral mediated expression of GCaMP in CA1, and implantation of a 1mm diameter GRIN lens just above the alveus (**Figure 3A**). Spatial memory was assessed using the DG-dependent spatial object recognition (SOR) task previously described [35]. During training, mice were exposed to an environment with three different objects then immediately given saline or 1 mg/kg CNO. Testing occurred 24 hours later when mice were reintroduced to the same box with one of the objects displaced (**Figure 3B**). Mice preferentially explore novelty and will spend more time around the displaced object resulting in a positive discrimination index (DI; see materials and methods for calculation). Mice were run through the SOR twice, receiving saline one time and 1 mg/kg CNO the other, with the order of treatment randomized across animals. Importantly, miniscope recordings were obtained approximately 1 hour after CNO injections to verify seizure activity. Because the goal was to investigate if induced seizures impaired spatial memory, animals that did not have seizures during this time were excluded from analysis. Animals successfully discriminated between the displaced and non-displaced object when they were given the saline injection (DI: 19.42 ± 33.63; mean ± StDev) but were impaired when they received 1 mg/kg CNO and had at least one verified seizure (DI: -6.295 ± 28.97; mean ± StDev; paired t test p = 0.0083; **Figure 3C**).

**Figure 3:**
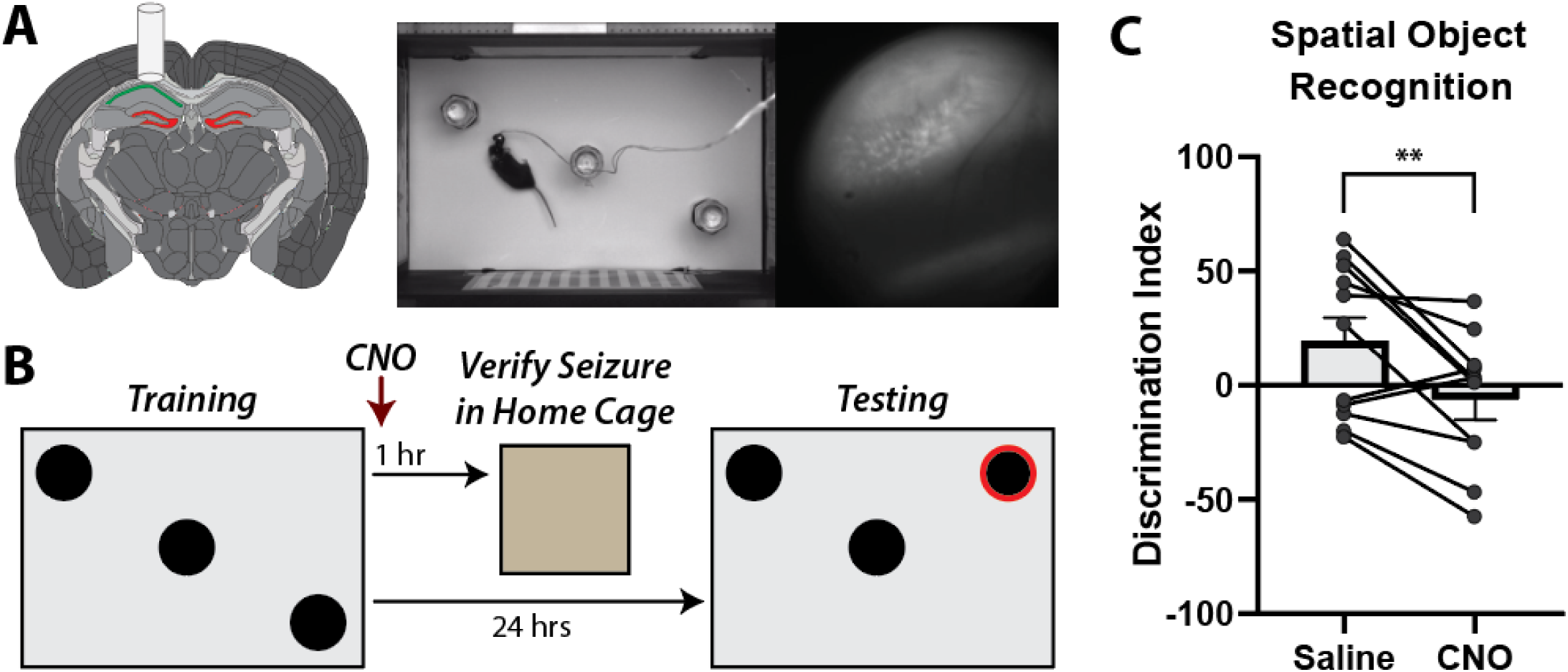
Induced seizures disrupt spatial memory in the SOR task. **A:** Schematic of the surgery with hM3Dq-mCherry expression in DGCs and GCaMP expression in CA1. Screenshot of a mouse in the SOR box with concurrent miniscope imaging. **B:** Schematic of the SOR paradigm. Mice were trained then immediately injected with 1 mg/kg CNO. One hour later, mice were recorded in their home cage for up to 30 mins to confirm seizure activity. Mice were then tested 24 hours later. **C:** SOR discrimination indices decrease after CNO injection compared to control saline injection (Saline: 19.42 ± 10.14; CNO: -6.30 ± 8.73; mean ± SEM; n = 11; paired t test; p = 0.0083).

### Seizures can be readily identified in miniscope recordings

During seizures, stereotypical patterns were observed in the calcium activity. In the first phase, the entire field of view would flash and become very bright. This was almost always followed by a spreading wave during which a small bright area would expand across the field of view. This was followed by a slow decrease in fluorescence across the entire field of view. Both the flashing and spreading wave were obvious during recording and can be seen when plotting the summed fluorescence of all pixels from the miniscope field of view (**Figure 4**). This activity pattern was observed when imaging from CA1 in the animals that were run through the SOR. Additionally, the same pattern of flashing followed by a spreading wave was observed in a separate cohort of mice when imaging from DGCs using a cre-dependent GCaMP and localizing the GRIN lens to the hippocampal fissure just at the edge of the outer molecular layer. Interestingly, this stereotypical activity pattern during seizures occurred in both CA1 and DG even when individual cells were not visible in the field of view. This is likely mediated by synchronous dendritic calcium spikes and/or out of focus cells.

**Figure 4:**
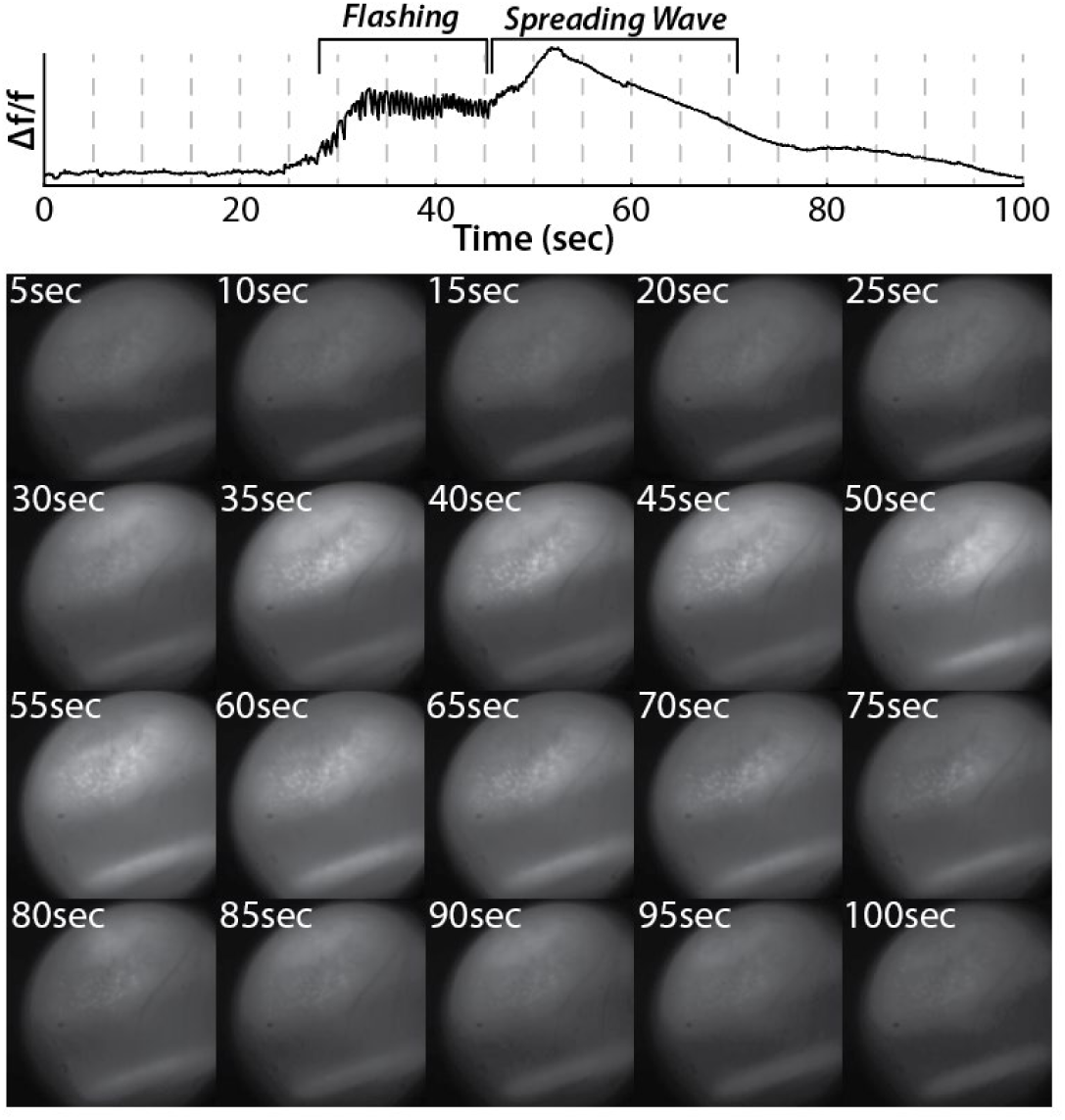
Seizures were reliably identified in miniscope imaging. **Top:** Δf/f from the entire miniscope field of view shows an epoch of flashing followed by a spreading wave. **Bottom:** Snapshots of miniscope frames matching dashed lines from Δf/f trace above.

## DISCUSSION & FUTURE DIRECTIONS

### Summary of findings

Using a targeted chemogenetic approach, we demonstrated that increasing excitability specifically in DGCs can cause seizures in non-epileptic mice (**Figures 1&2**). These results support the dentate gate hypothesis because DREADD-mediated increased excitability was restricted to DGCs which have been shown to be hyperexcitable in TLE [31, 35]. Importantly, induced seizures in this study were driven by intrinsic circuit activity rather than direct activation of DGCs previously reported [38]. This demonstrates that normal physiological inputs can drive seizures when DGCs are made to be hyperexcitable. Additionally, we show a dose-dependence of seizure severity which suggests that dentate gate degradation may be graded. This is further supported by a study showing that repeated optogenetic stimulations of DGCs worsened seizure severity [38]. Importantly, we also demonstrated that DGC-driven seizures can disrupt spatial memory in the DG-dependent SOR task when induced after training (**Figure 3**).

### Disentangling seizures and TLE pathologies

Deficits in hippocampal-dependent learning and memory tasks have been well documented in TLE [19, 35, 36, 41], and there is evidence that circuit manipulations in the hippocampus can rescue these behavioral deficits in animal models [35, 36]. One study showed that chemogenetically decreasing DG excitability in epileptic mice during training and testing could rescue behavioral deficits in the SOR task [35]. Another study found that shortening seizures for 2 weeks before behavior improved performance in the SOR task [36]. These studies highlight two distinct mechanisms that may disrupt behavior in TLE: i) pathological changes to hippocampal circuits and ii) seizure specific effects. These two mechanisms are difficult to untangle as hippocampal circuit disruptions can lead to seizures, and seizures can worsen hippocampal pathologies. The Kahn et al study addressed this problem by specifically testing whether pathological circuit changes affected spatial memory in the absence of seizures. By expressing hM3Dq in both DGCs and mossy cells, they were able to show that increasing DG excitability before testing in the SOR resulted in a spatial memory deficit that mirrored deficits in epileptic animals. Video-EEG monitoring of control animals was performed to demonstrate that no seizures were elicited following CNO injections at the does used for behavior. To further control for seizure effects, epileptic animals in this study were monitored during and proximal to behavior and were excluded if behavioral seizures were observed. The Kim et al study showed that shortening seizures rescued SOR behavioral deficits in epileptic mice; however, it could not disentangle whether effects were due to differences in the severity of seizure-drive hippocampal circuit disruptions, or whether effects were directly related to seizure activity. While both of these mechanisms are likely involved, our study provides direct evidence that DGC driven seizures can disrupt behavior in the absence of pathological changes to hippocampal circuitry.

Other studies have tried to disentangle the effects of pathological circuit disruptions and seizures on learning and memory deficits. Studies using flurothyl to induce seizures in non-epileptic animals showed that spatial memory could be transiently disrupted following a single seizure or multiple days of induced seizures, but performance recovered over time [42–44]. Behavioral performance and place cell stability was disrupted 20 minutes after a single flurothyl-induced seizure but returned to baseline 24 hours later [43]. Additionally, behavioral deficits observed following daily flurothyl-induced seizures did not occur for the first few days of induced seizures [42, 44]. While these studies appear to contradict our results, it is important to note that in these experiments, animals had already gone through multiple days of training before seizures were induced. This suggests that previously consolidated memories may be less susceptible to seizure-induced disruptions. Supporting this idea, one study found that inducing a seizure with PTZ after the critical learning day of a T-maze task impaired testing 24 hours later [37]. They also reported increased plasticity and saturated LTP in seizure tagged neurons as well as an overlap in the population of CA1 pyramidal cells activated in the T maze task and following the induced seizure. This suggests that cells participating in behavior may also be recruited to fire during seizures resulting in aberrant activity, and that plasticity may “overwrite” recently encoded events. This is further supported by our finding that DGC-driven seizures disrupt spatial memory in the DG-dependent SOR task when induced after training.

### Potential mechanisms underlying behavioral deficits

DGCs are thought to be important for memory encoding and may potentiate entorhinal inputs to CA3 as well as CA3 recurrent collaterals through heterosynaptic plasticity [45]. Through this mechanism, DGCs aberrantly activated during seizures could change synaptic weights in CA3 effectively “overwriting” recently encoded representations stored there resulting in retrograde amnesia. It has been shown that DGCs active during behavior have a transient increase in excitability [46], so it likely that these cells would be activated again when seizures occur soon after behavior. These cells may even be preferentially reactivated as the intrinsic excitability of DGCs is especially important for determining whether or not they will fire during behavior [47, 48]. Importantly, one study investigated whether there was a regional-specific overlap in populations of cells active both during a T-maze task and following a PTZ-induced seizure. They found an overlap in the populations in the retrosplenial cortex and medial prefrontal cortex, but not in the DG [49]. However, behavioral tagging and seizure tagging occurred 6 days apart which may explain the lack of overlap in DG as DGCs have increased intrinsic excitability after learning that is transient [46], and the sparse firing of DGCs and generally low excitability biases them to not fire across different conditions [47]. Additionally, it is important to interpret activity dependent labeling studies carefully as different techniques have different sensitivities, and labeling can underestimate actual activity. While the precise mechanism underlying behavioral deficits following induced seizures remains unknow, future studies that record activity during training, induced seizures, and testing may provide some insight.

### Implications and future directions

Our DGC-specific, chemogenetically induced seizure strategy is ideal for investigating mechanisms underlying behavioral deficits that occur following seizures. The known seizure focus in DG is relevant for TLE as the dentate has been shown to be hyperexcitable in TLE [31–35, 40]. This is preferable to flurothyl or PTZ induced seizures which do not have a known seizure focus. Additionally, we have confirmed that DGC-induced seizures can be readily identified in calcium imaging [50] (**Figure 4**), and miniscope imaging can occur during training and testing in the SOR. This allows network-level mechanisms underlying behavioral deficits to be investigated. For example, there is evidence that place cell stability may be disrupted following induced seizures [43, 44] which could be directly investigated through miniscope imaging. Future studies could also leverage miniscope imaging to explore whether task specific cells, such as object-vector cells in the SOR or splitter cell in the T-maze, are disrupted following DGC-induced seizures.

Additionally, the dose dependent severity of DGC induced seizures could be leveraged to investigate how small focal hippocampal seizures and interictal spiking may disrupt behavior and hippocampal coding. There is evidence that interictal spiking and high frequency oscillations may disrupt hippocampal coding in epilepsy [51–57], but the effects of these events have not been investigated in non-epileptic animals. Using our chemogenetic strategy, a low dose of CNO (0.3 mg/kg) can elicit interictal spikes which allows network level effects of these events to be further investigated. Doing this in non-epileptic animals removes any confounds from pathological changes to circuits that occur in epilepsy. Additionally, evoked events are transient and anchored to an initiating event meaning that events can be elicited at specific times such as before, during, or after behavior. Future studies could investigate whether interictal spiking at any of these specific times preferentially disrupts behavior. Additionally, the activity of individual cells could be tracked during interictal spikes or throughout the process of secondary generalization of hippocampal seizures to better understand the micro macro disconnect between the activity of individual cells and network level seizures [58]. While recording the activity of individual cells is difficult during seizures, recording single cell activity with calcium imaging or electrophysiology before and after seizures may shed light onto seizure initiation, propagation, and termination mechanisms, especially when cell type specific recording strategies are used. In addition to better understanding the mechanisms and effects of seizures, DGC-induced seizures could be used to test interventions such as anti-epileptic drugs and neuromodulatory strategies to reduce seizure burden and rescue cognitive deficits. Our DGC chemogenetic seizures can be repeatably induced, have effects that are restricted in time, have dose-dependent severity, occur in the absence of pathological circuit changes in epilepsy, and have a known seizure focus in the DG implicating their relevance in TLE. Seizures induced using this strategy provide direct evidence for the dentate gate theory as increasing excitability specifically in DGCs caused seizures in non-epileptic animals and resulted in cognitive deficits.

## MATERIALS AND METHODS

### Animals

Animal care and all procedures in this study were approved by the Children’s Hospital of Philadelphia Institutional Animal Care and Use Committee. Rbp4-cre^+^ (RRID:MMRRC_037128-UCD) mice bred with wild-type C57BL/6 mice (Charles River Laboratories) were used for all experiments. Mice were kept on a 12 hour light/dark cycle and had *ad libitum* access to food and water.

### Surgeries

#### Viral injection

Mice were anesthetized with isoflurane and affixed to a stereotaxic holder. A midline incision was then made on the head and small burr holes were drilled bilaterally into the skull (AP:-2.1 mm ML:1.2 mm). A needle was slowly advanced into each burr hole until the tip reached the dentate gyrus (AP:-2.1 mm ML:1.2 mm DV:2.0 mm), then 200nL of virus (AAV5-hSyn-DIO-hM3Dq-mCherry; Addgene 44361-AAV5) was injected at 100nL/min and the needle was left in place for 5 minutes before slowly retracting out of the brain. The midline incision was then sutured, and mice received buprenorphine E.R. (0.5 mg/kg s.c.) for pain management.

#### EEG implantation

Mice were anesthetized with isoflurane and affixed to a stereotaxic holder. The midline incision from the viral injection surgery was then reopened to visualize the skull. Stainless steel screws used for ground and reference were placed posterior to the lambda skull suture on either side of midline with their tips touching the brain. An additional two screws were used as cortical electrodes and were placed anterior to the bregma skull suture on either side of midline with their tips touching the brain. The existing burr hole on the right side was widened and a hippocampal depth electrode consisting of 2 stainless steel wires twisted together was inserted into the dorsal hippocampus targeting the CA1 region (AP:-2.1 mm ML:1.3 mm DV:1.2 mm). See **Figure 1A** for recording setup. Electrodes were held in a six-pin pedestal and were secured with dental cement. Mice were given buprenorphine E.R. (0.5 mg/kg s.c.) for pain management and allowed to recover for one week before undergoing video-EEG monitoring.

#### GRIN lens implantation

Mice were anesthetized with isoflurane and affixed to a stereotaxic holder. The midline incision from the viral injection surgery was then reopened to visualize the skull. A circular craniectomy (∼2.5 mm diameter) was made over the existing burr hole on the left side. Cold sterile saline was perfused onto the brain, and cortex underneath the craniectomy was gently aspirated until the corpus collosum became visible. Fibers of the corpus collosum were then carefully aspirated until the alveus became visible. After stabilizing all bleeding, a 1 mm GRIN lens (Inscopix) was placed on top of the alveus (AP:-2.1 mm ML: 1.4 mm) and secured with cyanoacrylate glue and dental cement. The lens was then covered in Kwik-Cast silicone sealant for protection, and mice were given buprenorphine E.R. (0.5 mg/kg s.c.) for pain management. GRIN lenses were coated in a mixture of GCaMP virus (AAV9-CaMKII-GCaMP6f-WPRE; addgene 100834-AAV9; or AAV9-syn-jGCaMP7f-WPRE addgene 104488-AAV9) and silk fibroin (Advanced Biomatrix 515420ML; 1 µL of 1:1 mixture). In a subset of mice, 300nL of GCaMP virus was stereotaxically injected into CA1 (AP:-2.1 mm ML:1.4 mm DV:1.5 mm) at least 20 minutes before beginning to aspirate cortex. In the cohort of DGC imaging mice, lenses were coated with a cre-dependent GCaMP (AAV9-syn-FLEX-jGCaMP8m-WPRE addgene 162378-AAV9) mixed 1:1 with silk fibroin and targeted to the hippocampal fissure just above outer molecular layer. This was achieved by aspirating to the alveus as described above then stereotaxically lowering a 1 mm biopsy punch into the aspirated area down to the hippocampal fissure. Tissue within the 1 mm diameter of the punch could be easily visualized and aspirated. GRIN lenses were then placed at the bottom of the aspirated area. At least 2 weeks after GRIN lens implantation, mice underwent a baseplating procedure. Mice were anesthetized with isoflurane and affixed to a stereotax holder. A miniscope attached to a baseplate was then lowered on a stereotax arm until a clear field of view was visualized in the miniscope recording software. Dental cement was applied to the miniscope baseplate, then the miniscope was removed and replaced with a protective cover for the lens.

### Behavior

#### Video-EEG recordings

Mice were continuously recorded using a Stellate-Harmonie (Stellate Inc., Montreal, Canada) 16-bit, 32 channel digital Video-EEG recording system. EEG recordings were sampled at 2 kHz and were time synchronized with video recordings. Mice were recorded for at least 1 week to establish a baseline and confirm that no seizures occurred spontaneously in implanted animals. Mice then received an i.p. saline injection followed by 4 different doses of CNO (0.1, 0.3, 1, and 3 mg/kg) injected i.p. and were video-EEG monitored to investigate the effects of these treatments on epileptic activity. Doses of CNO were administered in a random order, and all injections were followed by a minimum 48-hour washout period except for the 3 mg/kg CNO dose which was followed by a minimum 72-hour washout. Secondary generalized seizure events were manually notated *post hoc*.

#### Spatial object recognition task

Mice were habituated to miniscopes by hooking them up and recording for at least 3 minutes in their home cage for a minimum of 5 days before starting behavior. The SOR task was run as described previously [35]. On training day, the miniscope was hooked up, and mice were recorded for 3 minutes in their home cage. Mice were then placed into an empty 50 x 30 x 30 cm box with distinct visual cues on each of the 4 sides for 6 minutes. Afterwards, mice were returned to their home cage for an additional 3 minutes. Three identical objects were put into the box in a specific configuration and mice were again placed into the box for 3, 6-min trials with 3-min home cage exposures following each trial. On the testing day (24-hrs later), the spatial configuration of the objects was changed by displacing one of the objects. Mice were again hooked up and recorded for 3 minutes in their home cage followed by 6 minutes in the newly configured box then by another 3 minutes in the home cage. Mice were run through the SOR paradigm twice with different objects used for each experiment and a minimum of 2 weeks between repeated testing. In one SOR test, seizures were induced by injecting CNO (1 mg/kg s.c.) immediately after testing. Approximately 1 hour later, miniscopes were again placed on the mice and recordings were obtained for up to 30 mins while mice were in their home cage. During this time, mice were monitored for seizures and any animals that did not have at least one confirmed seizure were excluded from behavioral analysis. Behavioral results were compared to control experiments in which no seizures were induced, but mice received a saline injection immediately after training. The order of objects used, object configurations, and injection order (saline vs CNO first) was randomized. Behavior videos were run through DeepLabCut [59] to obtain tracking. Tracking was cleaned by removing all points with a likelihood below 95%, interpolating removed points, then smoothing using a gaussian window with a width of 0.25 sec. Object interactions were computed by calculating the amount of time the animal’s head was within a specific radius of each object using a custom MATLAB script. Behavioral performance was quantified by calculating a discrimination index as follows:

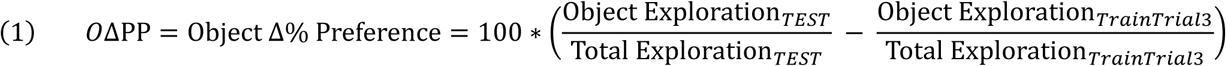

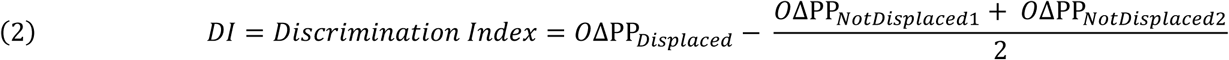

## AUTHOR CONTRIBUTIONS

Conceptualization: CDA, DAC

Formal Analysis: CDA, SMJ, HT

Investigation: CDA, ES, AW, SMJ

Resources: DAC

Writing – Original Draft Preparation: CDA

Writing – Review & Editing: CDA, SMJ, HT, DAC

Visualization: CDA

Supervision: DAC, HT

Funding Acquisition: DAC

## ACKNOWLEDGMENTS

This work was supported by funding from NIH NINDS R01NS038572 (DAC) and R01NS082046 (DAC).

